# SUMO-2/3 modification of PINK1 restrains basal mitophagy through MAPL-dependent regulation of mitochondrial surveillance

**DOI:** 10.1101/2025.05.27.656300

**Authors:** Nitheyaa Shree Ramesh, Richard Seager, Kevin A. Wilkinson, Jeremy M. Henley

**Affiliations:** Centre for Synaptic Plasticity, School of Biochemistry, Biomedical Sciences Building, University of Bristol, University Walk, Bristol, BS8 1TD, UK; Centre for Synaptic Plasticity, School of Physiology, Pharmacology and Neuroscience, Biomedical Sciences Building, University of Bristol, University Walk, Bristol, BS8 1TD, UK

**Author notes:** Max Planck Florida Institute for Neuroscience (MPFI), One Max Planck Way, Jupiter, Florida, 33458 USA.

## Abstract

The mitochondrial kinase PTEN-induced kinase 1 (PINK1) initiates mitophagy in response to mitochondrial damage, yet mechanisms that constrain its activation under basal conditions remain incompletely defined. Here, we identify a non-canonical SUMO-2/3 modification of PINK1 that suppresses basal mitophagy. We show that the mitochondrial E3 ligase mitochondrial-anchored protein ligase (MAPL/MUL1) promotes PINK1 SUMO-2/3 modification, while MAPL depletion reduces SUMOylation and enhances basal mitophagy. Targeted deSUMOylation increases PINK1 stabilisation and augments mitophagy. These findings reveal a novel SUMO-based checkpoint for basal PINK1 activation and mitochondrial quality control.

## Introduction

Mitophagy, the removal of damaged mitochondria, is fundamental for the maintenance of cellular health and function (Youle & Narendra, 2011), and its dysregulation is a prominent feature of a wide range of diseases, including neurodegenerative disorders, cancer, liver disease, and cardiovascular disease (Wang et al., 2023).

PTEN-induced kinase 1 (PINK1) is an essential sensor for the initial detection and subsequent clearance of damaged mitochondria (Geisler et al., 2010; Matsuda et al., 2010; Narendra et al., 2010; Vives-Bauza et al., 2010). Mutations in PINK1 cause autosomal recessive early-onset Parkinson’s disease, emphasising the essential role of PINK1-dependent mitophagy in maintaining cellular health (Gonçalves & Morais, 2021; Valente et al., 2004).

The stress-induced PINK1/Parkin mitophagy pathway has been extensively studied (for a recent review see (Narendra & Youle, 2024)). Briefly, PINK1 accumulates on the outer mitochondrial membrane (OMM) in response to membrane depolarisation (Geisler et al., 2010; Matsuda et al., 2010; Narendra et al., 2010). It then autophosphorylates to activate its kinase domain, leading to the phosphorylation and subsequent activation of the cytosolic E3 ubiquitin ligase, Parkin (Okatsu et al., 2012; Zhuang et al., 2016). PINK1 also directly phosphorylates ubiquitin at serine 65, which promotes Parkin recruitment to the mitochondria (Kane et al., 2014; Kazlauskaite et al., 2014; Koyano et al., 2014). Phosphorylated Parkin then ubiquitinates other OMM proteins. In a positive feedback loop, ubiquitinated OMM proteins recruit more Parkin to ubiquitinate more OMM proteins (Ordureau et al., 2014). This accumulation of ubiquitinated OMM proteins engages adaptor proteins such as p62, NDP52, and OPTN, which bind to LC3-II receptors on the autophagosome membrane (Lazarou et al., 2015; Narendra et al., 2010). Although stress-induced PINK1-mediated mitophagy is well characterised, the role, if any, of PINK1 in regulating basal mitophagy remains highly controversial (Ganley et al., 2021; Y.-T. Liu et al., 2021b, 2021a; McWilliams et al., 2018).

Under basal conditions, PINK1 is imported into the mitochondria, and subsequently cleaved by two proteases, mitochondrial protein peptidase (MPP) between residues P34 and G35 (Greene et al., 2012), and presenilin-associated rhomboid-like protein (PARL) between residues A103 and F104 (Deas et al., 2011; Jin et al., 2010). This double cleavage yields a mature ∼52 kDa form of PINK1, which is then retrotranslocated to the cytosol and targeted for proteasomal degradation (Yamano & Youle, 2013). Treatment of cells with the mitochondrial uncoupler, carbonyl cyanide m-chlorophenylhydrazone (CCCP), stabilises PINK1 at the mitochondrial membrane, while treatment of cells with the proteasomal inhibitor, MG132, inhibits degradation of the retrotranslocated cytosolic mature PINK1 (Narendra et al., 2010).

PINK1 contains multiple phosphorylation sites and, while the function of some remains unknown (Luo et al., 2024; Waddell et al., 2023), it has been established that autophosphorylation at S228 or S402 regulates the kinase activity of PINK1 (Aerts et al., 2015; Okatsu et al., 2012). In addition, S167 and C412 have also been identified recently as autophosphorylation sites affecting PINK1 activation (Luo et al., 2024; Waddell et al., 2023).

Ubiquitination of the 52 kDa form of PINK1 at K137 targets it for proteasomal degradation (Y. Liu et al., 2017). Additionally, PINK1 can undergo degradation via the N-end ubiquitination rule, independent of K137 ubiquitination (Yamano & Youle, 2013). In response to specific stressors, PINK1 can also be S-nitrosylated at C568, leading to a decrease in kinase activity and reduced mitochondrial translocation of Parkin (Oh et al., 2017).

SUMO-2/3-ylation is the covalent addition of a small ubiquitin-like modifier (SUMO2/3) protein to a substrate, canonically at a lysine residue within a SUMO consensus motif (Vertegaal, 2022; Wilkinson & Henley, 2010). SUMO conjugation plays critical roles in protein subcellular localisation, stability, and function (Henley et al., 2018; Wilkinson & Henley, 2010). Briefly, SUMO is a ∼11 kDa protein, of which there are three mammalian isoforms (SUMO1, SUMO2, and SUMO3). SUMO2 and SUMO3 differ by only three amino acid residues (∼97% identity) and are generally referred to collectively as SUMO2/3 (Saitoh & Hinchey, 2000). SUMO1 shares only ∼50% identity with SUMO2/3. Both SUMO1-ylation and SUMO2/3-ylation are highly dynamic, reversible post-translational modifications with conjugation mediated by an E1, E2, and E3 enzyme cascade and deconjugation mediated by specific SUMO proteases, most notably SENPs (Celen & Sahin, 2020; Hay, 2005).

Here, we show that PINK1 can undergo SUMO-2/3-ylation by the SUMO E3 ligase MAPL. Remarkably, SUMO-2/3-ylation of PINK1 does not occur at lysine residues, since a mutant PINK1 in which all lysines were substituted with arginines is still SUMO-2/3-ylated. Nonetheless, by perturbing SUMO-2/3-ylation of PINK1, we demonstrate that SUMO-2/3-ylation of PINK1 mediates PINK1-dependent mitophagy. Together, these findings indicate that non-canonical MAPL-mediated SUMO-2/3-ylation of PINK1 contributes to regulatory mitochondrial surveillance in health and disease.

## Results

### PINK1 is modified by SUMO2/3

To examine potential SUMO-2/3-ylation of PINK1, PINK1-GFP was overexpressed in HEK293T cells. We used 2% SDS denaturing conditions in PINK1-GFP immunoprecipitation experiments to disrupt non-covalent interactions but conserve post-translational modifications. In immunoblots with an anti-SUMO2/3 antibody we observed a higher molecular weight ladder in the PINK1-GFP pull-downs, indicative of poly-SUMO chains (Fig. 1A). We also immunoblotted the PINK1-GFP immunoprecipitate for ubiquitin. Similar to SUMO2/3, we also observed a ladder indicative of PINK1-GFP polyubiquitination (Fig. 1A). Co-transfecting the SBP-tagged SUMO protease, SBP-SENP1, significantly decreased the SUMO2/3 smear on PINK1-GFP (Fig. 1B), but did not alter the ubiquitin smear. Interestingly, we also observed that the total amount of PINK1-GFP in cells increased significantly upon co-expression of SBP-SENP1 (Fig. 1C).

**Figure 1.**
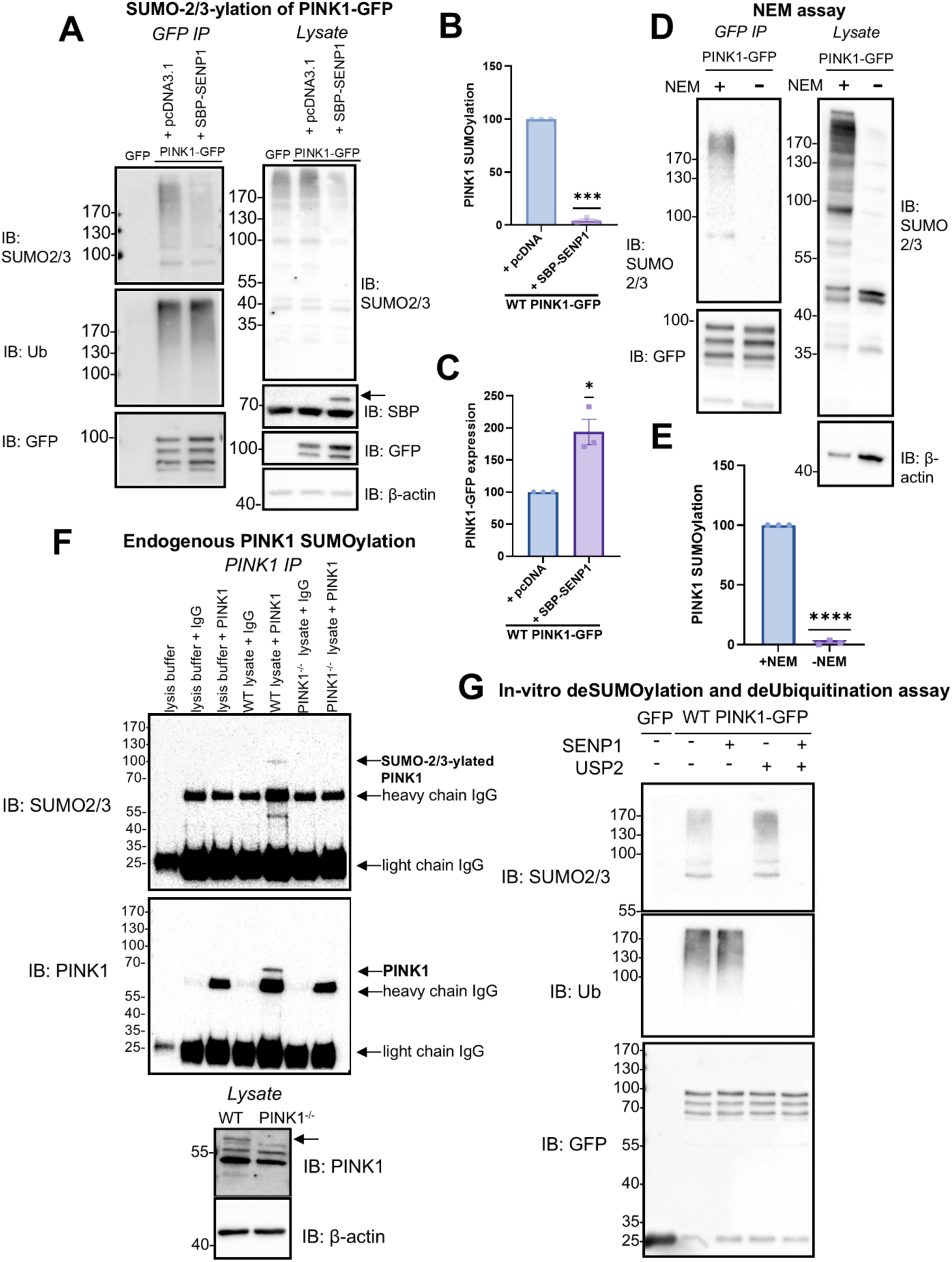
PINK1 is SUMO-2/3-ylated. **(A)** HEK293T cells were transfected with GFP, PINK1-GFP, and pcDNA3.1 or SBP-SENP1. Representative blot of SUMO2/3 and Ub higher molecular weight smears on immunoprecipitated PINK1-GFP and corresponding input samples with full-length PINK1-GFP (FL) and cleaved PINK1-GFP (CL). **(B)** Quantification of SUMO2/3 smear on PINK1-GFP immunoprecipitation (SUMO2/3/GFP normalized to control (PINK1-GFP + pcDNA)) (n=3, one-sample t-test, ***p<0.001) **(C)** Quantification of total PINK1-GFP expressed in cells (GFP/actin normalized to control (PINK1-GFP + pcDNA)) (n=3, one-sample t-test, *p<0.05) **(D)** HEK293T cells overexpressing PINK1-GFP were lysed in the presence or absence of n-ethylmaleimide (NEM), a cysteine protease inhibitor. Representative blots of SUMO2/3 higher molecular weight smear on PINK1-GFP and corresponding inputs. **(E)** Quantification of SUMO2/3 smear on immunoprecipitated PINK1 (SUMO2/3/GFP normalized to control (+NEM)) (n=3, one-sample t-test, ****p<0.0001) **(F)** Endogenous PINK1 was immunoprecipitated from mitochondrial fractions prepared from WT and PINK1^-/-^ HEK293T cells, and samples were immunoblotted for SUMO2/3 and PINK1. 10% input of both WT and PINK1^-/-^ cells were immunoblotted for PINK1 and β-actin. **(G)** HEK293T cells were transfected with GFP or PINK1-GFP for 48 hours. Cells were lysed and PINK1-GFP was immunoprecipitated using GFP-trap beads. Active SUMO protease, SENP1 (100 nM) or active deubiquitinase, USP2 (500 nM) were added to IP samples for 2 hours at 37° C, samples were run on SDS-PAGE gel and immunoblotted for SUMO2/3 and ubiquitin.

To further corroborate that the PINK1 smear was due to SUMO2/3-ylation, the cysteine protease inhibitor n-ethylmaleimide (NEM), which inhibits the cysteine protease activity of the SENPs, was included in the lysis buffer before PINK1-GFP immunoprecipitation (Fig. 1D). The PINK1-SUMO2/3 smear was significantly increased in cells lysed in NEM (Fig. 1E). Taken together, these data indicate that exogenously expressed PINK1 is SUMO2/3-ylated.

We next immunoprecipitated endogenous PINK1 from HEK293T cell mitochondrial-enriched fraction using an anti-PINK1 antibody. When blotted for SUMO2/3 we detected a single band in the PINK1 pulldown lane that was not present in the mitochondrial-enriched fraction from PINK1^-/-^ HEK293T cells (Fig. 1F). Interestingly, in these endogenous PINK1 blots, only one clear band appears and overexposure of the blot does not reveal a distinct higher molecular weight smear, differing from overexpression experiments present in Fig. 1A. Additionally, this band is not present when immunoblotting for PINK1. We attribute these two observations to the very low levels of SUMO-2/3-ylated endogenous PINK1 in cells, especially given that there is little PINK1 in healthy cells (Narendra et al., 2010).

Since we observe both poly-SUMO-2/3-ylation and poly-ubiquitination on PINK1 (Fig. 1A), we wondered if these may be mixed SUMO-ubiquitin chains (Hendriks et al., 2014). To assess the composition of the higher molecular weight smear on PINK1-GFP, we applied constitutively active recombinant SENP1 or the deubiquitinating enzyme USP2 to immunoprecipitated PINK1-GFP from HEK293T cells (Fig. 1G). Active SENP1 removed SUMO2/3 from PINK1-GFP but did not affect ubiquitin, whereas active USP2 removed ubiquitin but did not affect SUMO2/3. Treatment with both active SENP1 and USP2 completely removed both ubiquitin and SUMO2/3 smears on PINK1-GFP (Fig. 1G). These data indicate that PINK1 can be simultaneously SUMO-2/3-ylated and ubiquitinated but that there are no mixed SUMO-ubiquitin chains.

### Lysine-independent PINK1 SUMO-2/3-ylation

Canonically, SUMO is conjugated to lysine residues. Therefore, to identify where PINK1 is SUMO-2/3-ylated, lysines in PINK1-GFP were systematically and additively mutated to non-SUMOylatable arginine residues. Initially, we mutated lysines previously identified to be ubiquitinated as well as lysines located within a SUMO consensus motif (ψ-K-x-D/E, where ψ is a hydrophobic residue) (Fig. 2A). Unexpectedly, none of these mutations individually or collectively altered PINK1-GFP SUMO-2/3-ylation. Indeed, even PINK1-GFP with all 21 lysines mutated to arginines (21KR PINK1-GFP) was still SUMO-2/3-ylated. Importantly, however, co-transfection with SENP1 dramatically decreased SUMO-2/3-ylation of both wild-type PINK1-GFP and 21KR PINK1-GFP (Fig. 2B). Notably, unlike levels of SUMO-2/3-ylation, levels of ubiquitin were significantly reduced 21KR PINK1-GFP (Fig. S2). These unexpected results demonstrate that PINK1 is reversibly SUMO2/3-ylated despite the removal of all lysines.

**Figure 2.**
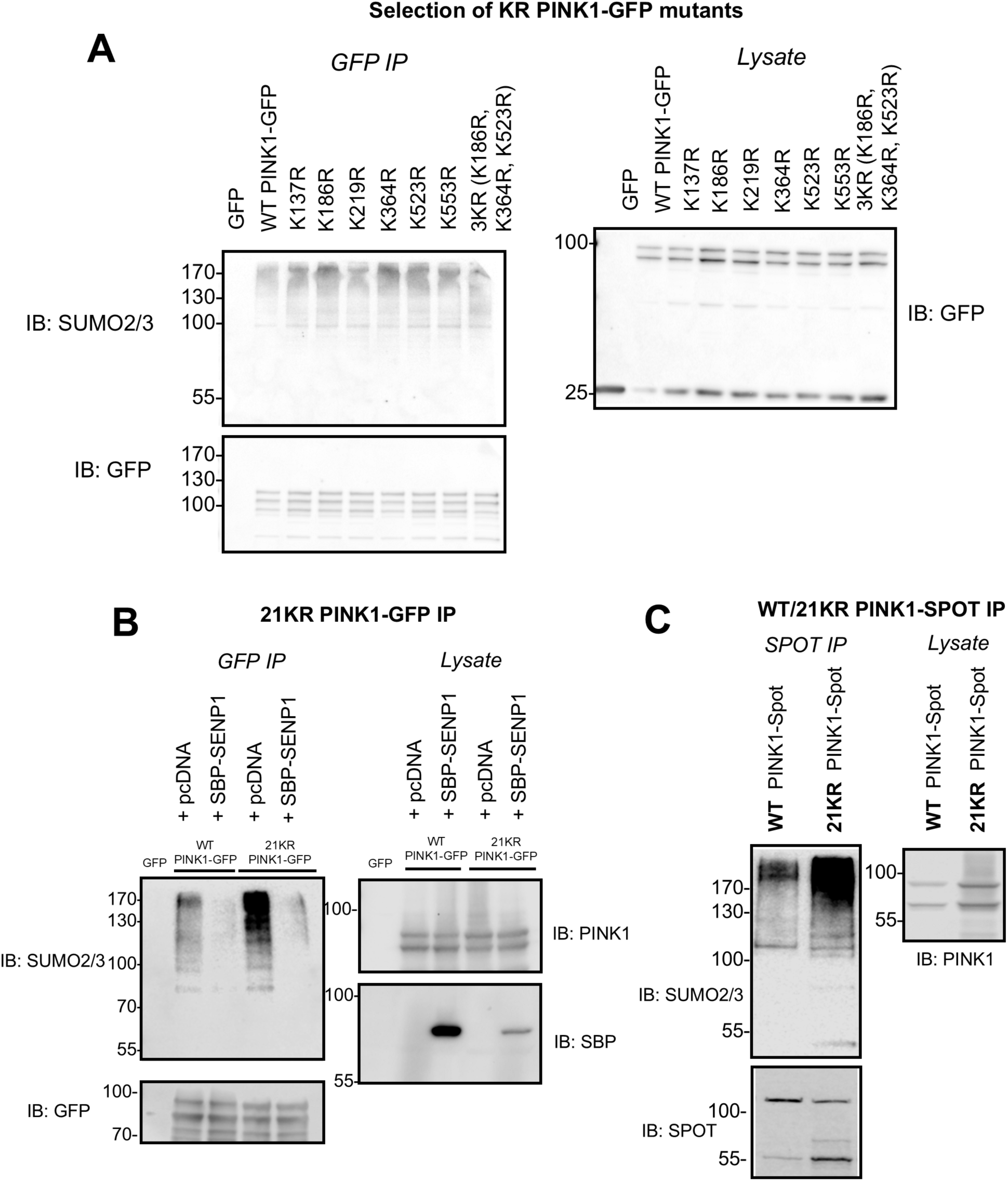
PINK1 is not SUMO-2/3-ylated at a lysine. **(A)** Initial set of PINK1-GFP lysine mutants. HEK293T cells were transfected with GFP, WT PINK1-GFP, or KR mutants of PINK1-GFP for 48 hours. PINK1 was immunoprecipitated using GFP trap beads, and samples were immunoblotted for SUMO2/3 and GFP. **(B)** HEK293T cells were transfected with WT PINK1-GFP or 21KR PINK1-GFP along with SBP-SENP1. GFP-Trap performed on lysate, and samples probed for SUMO2/3 and GFP. Lysates were probed for PINK1 and SBP. **(C)** WT and 21KR PINK1-Spot were overexpressed in HEK293T cells and PINK1 was immunoprecipitated using ChromoTek Spot-Trap beads. Samples were then immunoblotted for SUMO2/3, Spot, and PINK1.

Intriguingly, it has been reported recently that substrates can be SUMO-2/3-ylated at the amino group of their N-terminal, similar to the well-established N-terminal ubiquitination (El Motiam et al., 2025; Weng et al., 2023). To test if this happens for PINK1, we co-transfected cells with WT PINK1-GFP and N-α-acetyltransferase 60 (Naa60), which irreversibly acetylates protein N-termini, thereby preventing further posttranslational modifications. When wild-type PINK1-GFP or 21KR PINK1-GFP was overexpressed with FLAG-Naa60, levels of SUMO-2/3-ylated PINK1 were not decreased (Fig. S1).

To make sure it was PINK1 SUMO-2/3-ylation and not the GFP tag, we exchanged the GFP for a SPOT tag (amino acid sequence: PDRVRAVSHWSS), thereby removing all lysine residues from the entire expressed protein (21KR PINK1-SPOT). Remarkably, both WT and 21KR PINK1-SPOT were still SUMO-2/3-ylated (Fig. 2C) indicating PINK1 SUMO-2/3-ylation is non-canonical and lysine-independent.

### MAPL promotes PINK1 SUMO-2/3-ylation

We reasoned a prime candidate to mediate PINK1 SUMO-2/3-ylation could be the dual specificity SUMO and ubiquitin ligase mitochondrial anchored protein ligase (MAPL). We used a SMART pool of siRNA targeting human MAPL to knock down MAPL in HEK293T cells (Fig. 3A). MAPL knockdown significantly decreased levels of PINK1 SUMO-2/3-ylation (Fig. 3B-3C). We next tested if MAPL binds to PINK1 by immunoprecipitating overexpressed PINK1-GFP and blotting for MAPL. As shown in Fig. 3D, MAPL was present in the WT PINK1-GFP lane but not in the GFP control (Fig. 3D). Importantly, the OMM protein VDAC did not co-immunoprecipitate with PINK1, indicating a selective interaction between PINK1 and MAPL.

**Figure 3.**
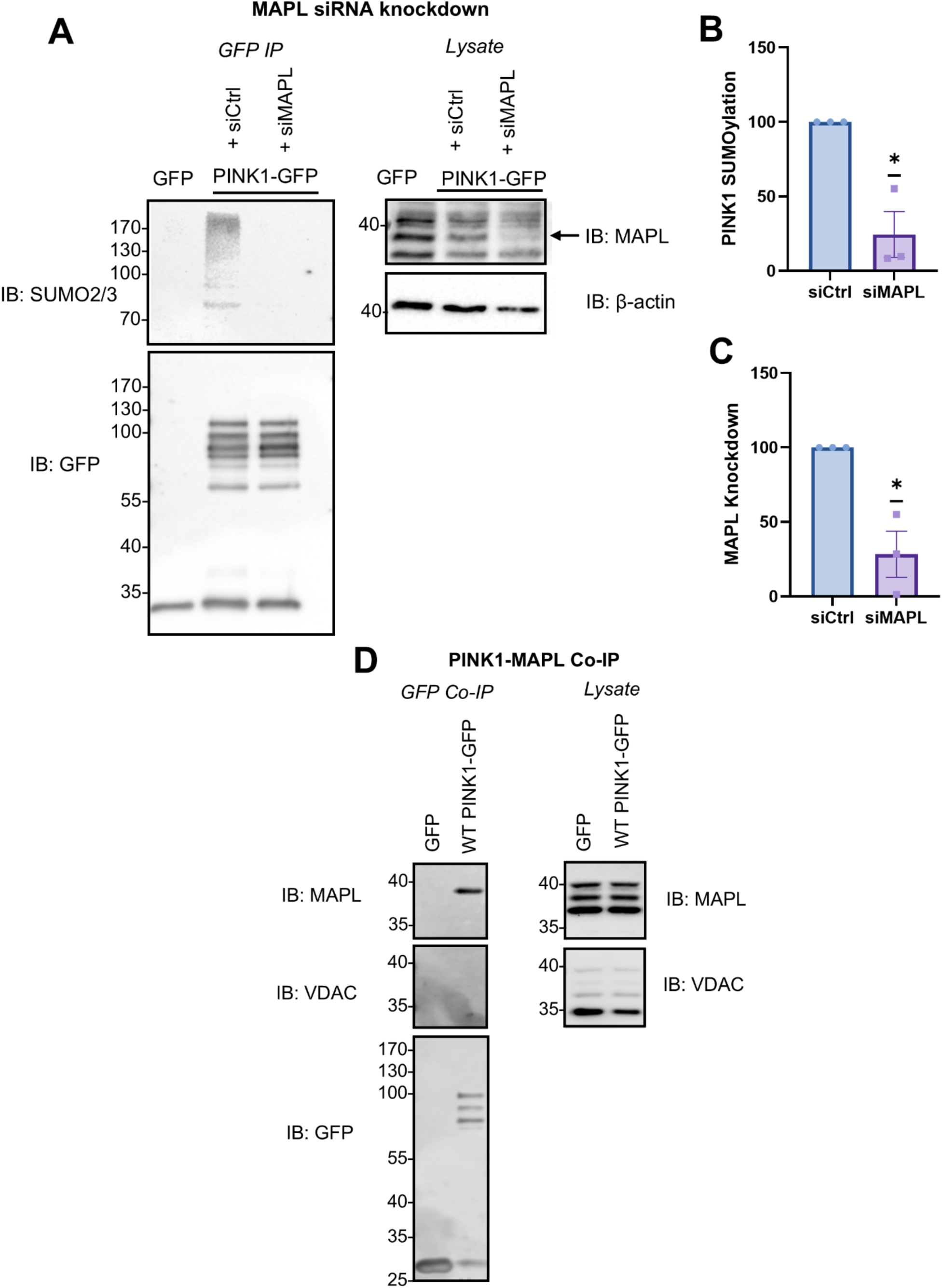
MAPL promotes PINK1 SUMO-2/3-ylation. **(A**) HEK293T cells were transfected with PINK-GFP and a control siRNA (20 nM) or MAPL siRNA (20 nM) for 48 hours. Representative blot of SUMO2/3 higher molecular weight smears on immunoprecipitated PINK1-GFP and corresponding input samples. **(B)** Quantification of SUMO2/3 smear on PINK1-GFP immunoprecipitation (SUMO2/3/GFP normalized to control (siCtrl)) (n=3, one-sample t-test, *p<0.05)**. (C)** Quantification of MAPL knockdown in total cell lysate (MAPL/actin normalized to control (siCtrl)) (n=3, one-sample t-test, *p<0.05) **(D)** HEK293T cells were transfected with WT PINK1-GFP and cells were lysed in buffer without SDS to preserve non-covalent interactions. PINK1 was immunoprecipitated using GFP trap beads and samples were immunoblotted for MAPL, VDAC, and GFP. Representative blot of at least three independent experiments.

### PINK1 SUMO-2/3-ylation and stability

PINK1 is constitutively degraded in healthy mitochondria with a reported half-life of ∼30 minutes (Lee et al., 2015; Lin & Kang, 2008). This constant turnover is mediated by mitochondrial import and cleavage of PINK1 (Deas et al., 2011; Greene et al., 2012). The remaining PINK1 fragment is translocated back to the cytosol where it is degraded via the proteasome (Jin et al., 2010). Conversely, in stressed mitochondria PINK1 is retained at the OMM thereby escaping MPP and PARL cleavage (Narendra et al., 2010).

To test the effects of SUMO2/3-ylation on PINK1 turnover, we carried out an array of experiments with CCCP and MG132, as well as two different point mutants of PINK1, P95A and F104A. CCCP dissipates the mitochondrial membrane potential and inhibits oxidative phosphorylation, which stabilises full-length PINK1 at the OMM (Narendra et al., 2010), whereas the proteasomal inhibitor MG132 accumulates the cleaved form of PINK1 (Lin & Kang, 2008). The P95A PINK1 mutant accumulates full-length PINK1 at the OMM, while the F104A mutant accumulates the cytosolic fragment of PINK1 (Deas et al., 2011).

In HEK293T cells, CCCP significantly decreased (Fig. 4A-B), whereas MG132 significantly increased (Fig. 4C-D), levels of SUMO2/3-ylation of PINK1. SUMO2/3-ylation of P95A PINK1-GFP, but not F104A PINK1-GFP, was reduced significantly compared to WT PINK1-GFP (Fig. 4E-F). Additionally, in total lysates from cells treated with CCCP there were higher levels of phospho-ubiquitin, reflecting full-length PINK1 activity (Fig. 4A). The level of total phospho-ubiquitin was also significantly higher for the P95A PINK1-GFP mutant compared to WT PINK1-GFP (Fig. 4G).

**Figure 4.**
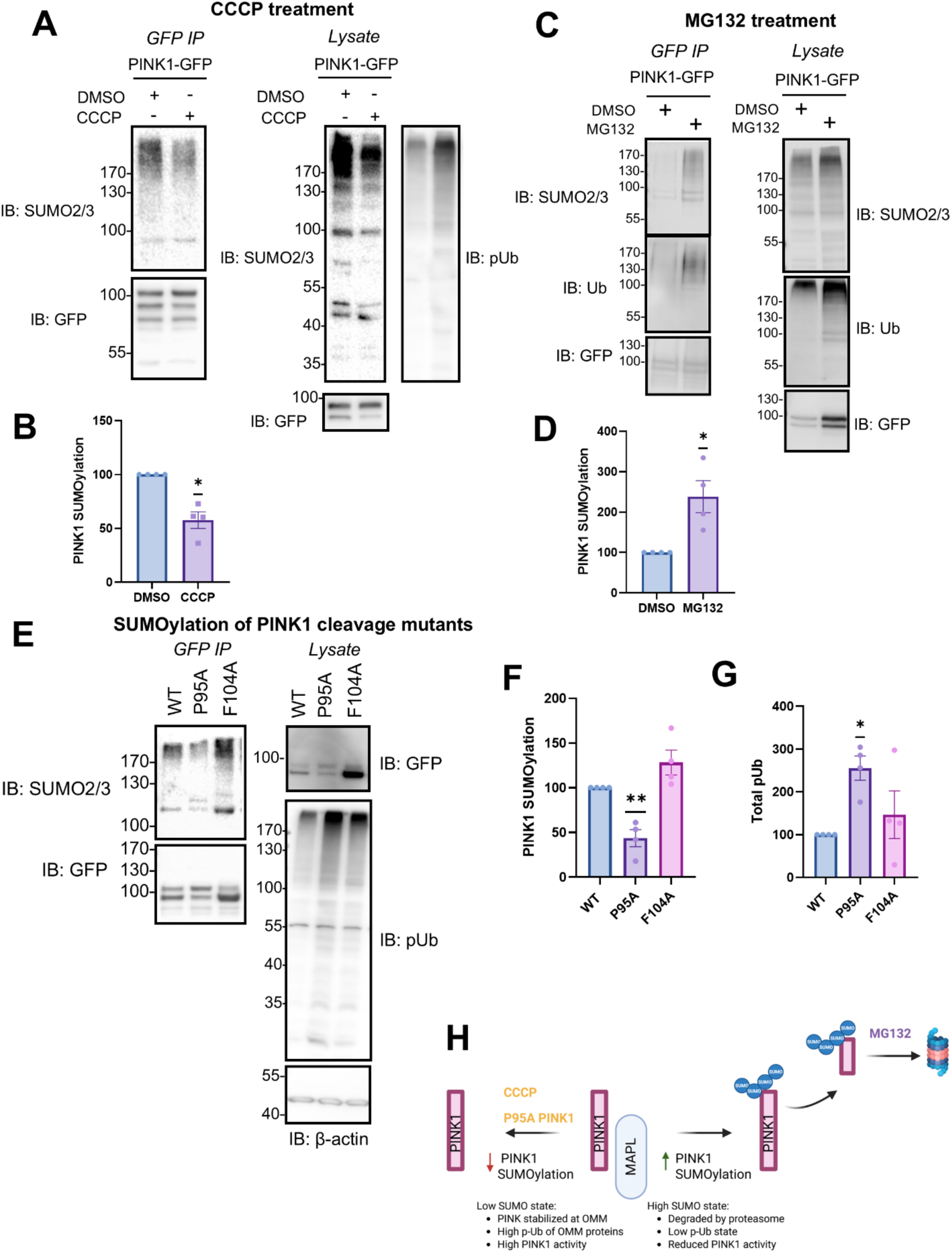
PINK1 SUMO-2/3-ylation and stability. **(A)** SUMO-2/3-ylation of PINK1 in response to CCCP treatment. HEK293T cells were transfected with PINK1-GFP and treated with 10 μM CCCP (or DMSO control) for one hour prior to lysis. Representative blots of higher molecular weight SUMO2/3 smear of immunoprecipitated PINK1 and corresponding input samples. **(B)** Quantification of higher molecular weight SUMO2/3 smear on PINK1 (SUMO2/3/GFP normalized to control (DMSO)) (n=4, one sample test *p<0.05) **(C)** SUMO-2/3-ylation of PINK1 in response to MG132 treatment. HEK293T cells were transfected with PINK1-GFP and treated with 20 μM MG132 (or DMSO control) overnight before immunoprecipitating PINK1 using GFP trap beads. Representative blots of higher molecular weight SUMO2/3 and ubiquitin smear on immunoprecipitated PINK1. **(D)** Quantification of SUMO-2/3-ylated PINK1 in DMSO and MG132 conditions (SUMO2/3/GFP normalized to control (DMSO)) (n=4, one sample t-test, *p<0.05) **(E)** HEK293T cells were transfected with WT, P95A, or F104A PINK1-GFP for 48 hours. Representative blots of higher molecular weight SUMO2/3 and ubiquitin smear on immunoprecipitated PINK1 along with corresponding inputs. **(F)** Quantification of higher molecular weight SUMO2/3 smear on PINK1 (SUMO2/3/GFP normalized to control (WT PINK1-GFP)) (n=4, one sample t-test, **p<0.01) **(G)** Quantification of total phosho-ubiquitin (S65) in lysates (pUb/GFP normalized to control (WT PINK1-GFP)) (n=4, one sample t-test, *p<0.05) **(H)** Potential model of the role of PINK1 SUMO-2/3-ylation in its stabilization: Both the P95A mutant of PINK1 and treatment of cells with CCCP decreased levels of SUMO-2/3-ylated PINK1 while treatment of cells with MG132 increased levels of SUMO-2/3-ylated PINK1. *Created in BioRender*.

In summary, CCCP treatment decreases levels of SUMO-2/3-ylated PINK1, while MG132 treatment increases levels of SUMO2/3-ylated PINK1, and the cleavage-resistant PINK1-P95A mutant exhibits reduced SUMO2/3-ylation (Fig. 4H). Taken together, these data suggest that non-SUMO2/3-ylated PINK1 is more stable.

### Counteracting PINK1 SUMO-2/3-ylation promotes mitophagy

We next tested if PINK1 SUMO-2/3-ylation affects PINK1-dependent mitophagy. WT and PINK1^-/-^ HEK293T cells were transfected with a control siRNA or siRNA targeting human MAPL for 48 hours and treated with 10μM CCCP (or DMSO control) for one hour. Samples were lysed and immunoblotted for a range of autophagy and mitophagy markers (Fig. 5A). MAPL knockdown increased the LC3-II/LC3-I ratio, indicative of increased autophagy (Fig. 5B). As expected, this increase was also observed in cells treated with CCCP. Importantly, MAPL-knockdown-mediated autophagy was absent in PINK1^-/-^ cells, confirming a PINK1-dependent pathway. Notably, however, CCCP increased the LC3-II/LC3-I ratio in PINK1^-/-^ cells, consistent with both PINK1-dependent and independent pathways, as previously reported (Seabright et al., 2020).

**Figure 5.**
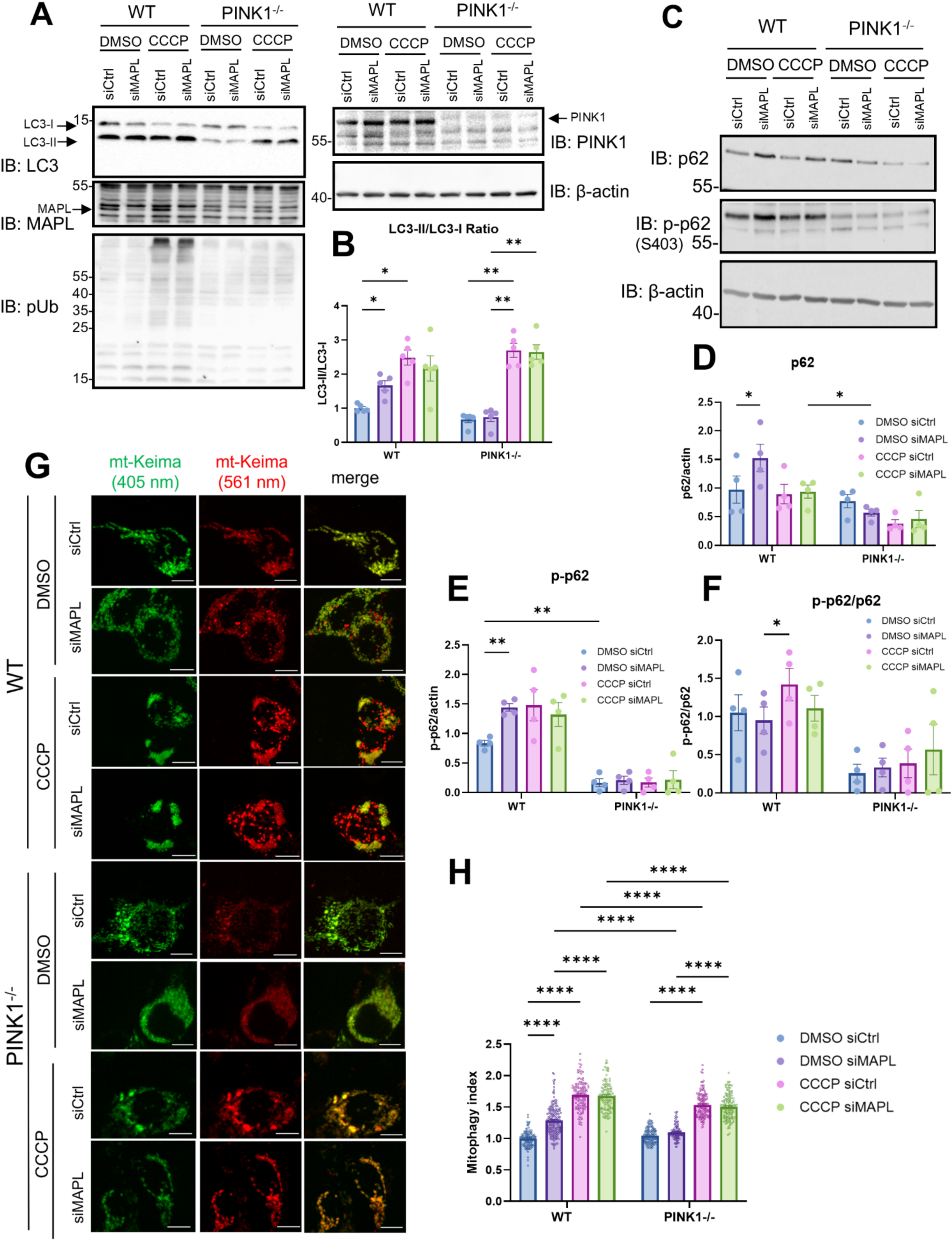
Knockdown of MAPL increases basal mitophagy. **(A and C)** Representative blots of WT and PINK1^-/-^ cells, transfected with a control or human MAPL targeting siRNA (20 nM). Cells were treated with 10 μM CCCP (or DMSO control) for one hour before lysis. Quantification of **(B)** LC3-II:LC3-I ratio (two-way ANOVA with Tukey’s post hoc test, n=5, *p<0.05, **p<0.01). Quantification of **(D)** p62, **(E)** phosphorylated p62 at S403, and **(F)** p-p62:p62 ratio (two-way ANOVA with Tukey’s post hoc test, n=4, *p<0.05, **p<0.01) **(G)** Representative images of WT and PINK1^-/-^ cells transfected with mitochondrial mtKeima (0.5 μg) and siRNA control (20 nM) or siRNA targeting human MAPL (20 nM) for 48 hours before imaging. Cells were treated with 10 μM CCCP (or DMSO control) for one hour before imaging. Scale bar denotes 10 μm. **(H)** Quantification of “mitophagy index”: fluorescence in 561 nm excitation channel divided by the total fluorescence in 405 nm and 561 nm excitation channels as a measure of “total mitophagy: total mitochondrial area” (C561/(C561 + C405)) (Kruskal-Wallis test followed by Dunn’s multiple comparison test, N=3, n= 101-189 cells, ****p<0.0001)

MAPL knockdown increased the ubiquitin-autophagy adaptor, p62 (SQSTM1), in WT but not in PINK1^-/-^ HEK293T cells (Fig. 5C-5D). Phosphorylated p62 was also increased by MAPL knockdown in WT cells, whereas levels of phosphorylated p62 were significantly lower in PINK1^-/-^ cells and were not further decreased by knockdown of MAPL (Fig. 5E-5F). These results suggest that MAPL knockdown, which strongly reduces PINK1 SUMO-2/3-ylation, enhances PINK1-dependent basal mitophagy.

We next visualised mitophagy using a mitochondria-targeted Keima-Red mitophagy probe. WT and PINK1^-/-^ HEK293T cells were transfected with mtKeima (pMitophagy Keima Red mPark2) and an siRNA targeting human MAPL (Fig. 5G). Consistent with our immunoblot data, MAPL knockdown increased mitophagy in WT but not in PINK1^-/-^ cells (Fig. 5H). Notably, MAPL knockdown did not cause a further increase in mitophagy in either WT or PINK1^-/-^ cells treated with CCCP for one hour, suggesting MAPL knockdown specifically affects basal mitophagy. Also consistent with the blot data, CCCP evoked mitophagy in both WT and PINK1^-/-^ cells, indicative of a PINK1-independent mitophagy pathway (Seabright et al., 2020) (Fig. 5H). Together, these results indicate that MAPL promotes basal mitophagy through its actions on PINK1.

Our data demonstrate that PINK1 is SUMO-2/3-ylated in a non-canonical lysine-independent manner. Consequently, despite extensive attempts, we were unable to identify a non-SUMOylatable mutant for loss-of-function studies. We therefore used an alternative approach to specifically and inducibly promote PINK1 deSUMO-2/3-ylation. We co-expressed PINK1-GFP with an anti-GFP-nanobody fused to the catalytic domain of SENP1 under a tetracycline promoter (GNb-SENP1) as described previously (Ibrahim et al., 2020; Mojsa et al., 2020). We reasoned doxycycline-induced GNb-SENP1 expression would selectively deSUMOylate PINK1-GFP (Fig. 6A). We validated this system by transfecting HEK293T cells with PINK1-GFP in the presence of the anti-GFP-nanobody (GNb), or the SENP1-fused anti-GFP-nanobody (GNb-SENP1) (Fig. 6B-6C). Doxycycline induction of GNb-SENP1 dramatically and selectively reduced PINK1 SUMO-2/3-ylation, with no evidence of differences in overall SUMO conjugation in the total lysate blots (Fig. 6D).

**Figure 6.**
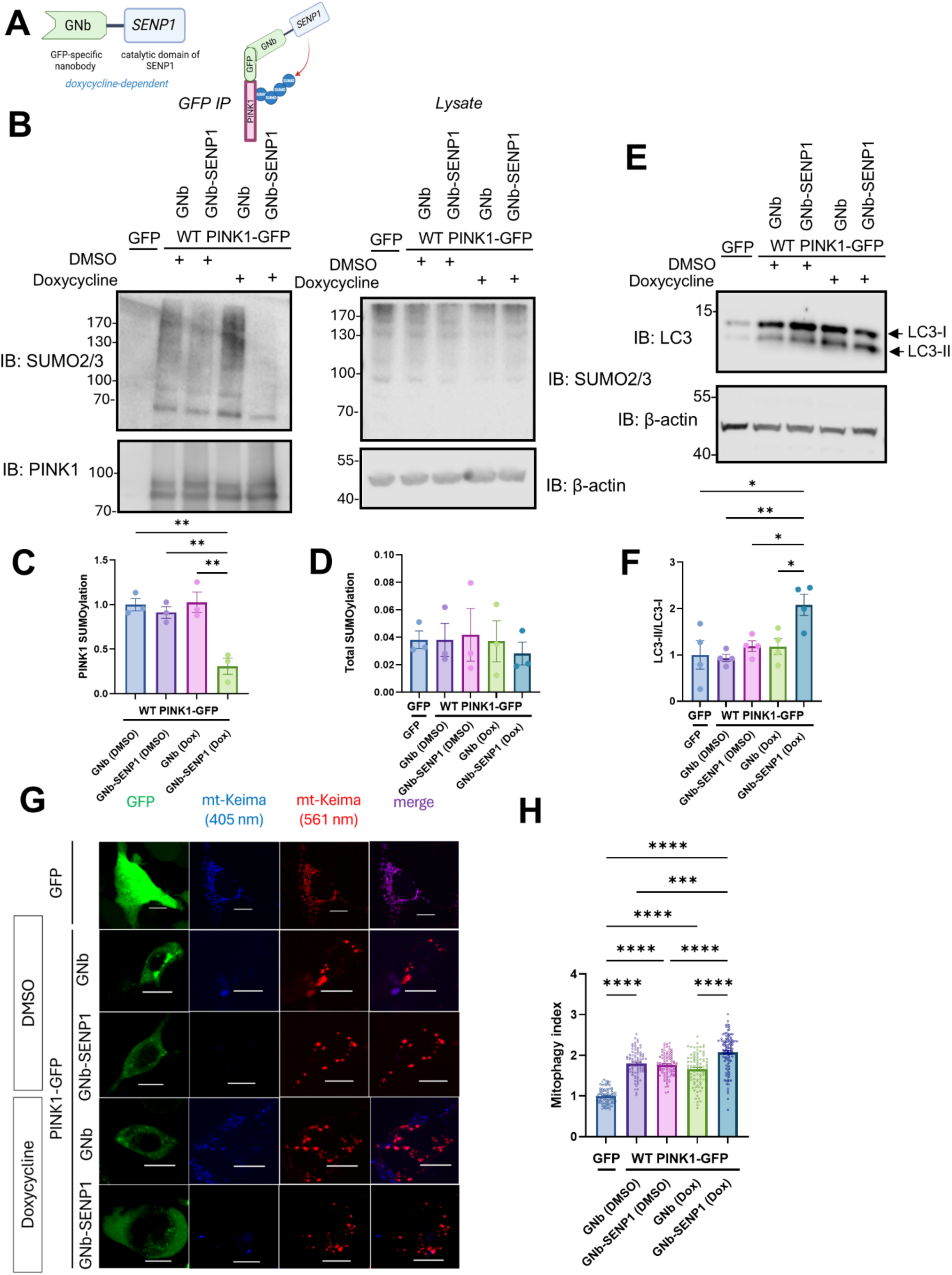
DeSUMO-2/3-ylation of PINK1 increases mitophagy. **(A)** Schematic of GNb-SENP1 deSUMOylating PINK1-GFP. **(B)** GNb-SENP1 selectively deSUMOylates PINK1. HEK293T cells were transfected with GFP or PINK1-GFP along with GNb or GNb-SENP1 for 72 hours. Cells were treated with doxycycline (or DMSO control) for 48 hours before immunoprecipitation. Representative blots of higher molecular weight SUMO2/3 smear of immunoprecipitated PINK1 and corresponding input samples. **(C)** Quantification of SUMO-2/3-ylated PINK1 (SUMO2/3/GFP) (n=3, one-way ANOVA with Tukey’s post-hoc test, **p<0.01) **(D)** Quantification of total SUMO2/3 in lysate (SUMO2/3/β-actin) (n=3, one-way ANOVA with Tukey’s post-hoc test) **(E)** Representative blots of total lysates from (B) immunoblotted for LC3A/B. **(F)** Quantification of LC3-II:LC3-I ratio (n=4, one-way ANOVA with Tukey’s post-hoc test, *p<0.05, **p<0.01) **(G)** Representative images of PINK1^-/-^ cells transfected with GFP or PINK1-GFP, mt-Keima, and GNb or GNb-SENP1. Cells were treated with 1 μg/mL doxycycline (or DMSO control) for 48 hours before imaging. Scale bar denotes 10 μm. **(H)** Quantification of “mitophagy index”: fluorescence in 561 nm excitation channel divided by the total fluorescence in 405 nm and 561 nm excitation channels as a measure of “total mitophagy: total mitochondrial area” (C561/(C561 + C405)) (Kruskal-Wallis test followed by Dunn’s multiple comparison test, N=3, n= 90-115 cells, ***p<0.001, ****p<0.0001)

**Figure 7.**
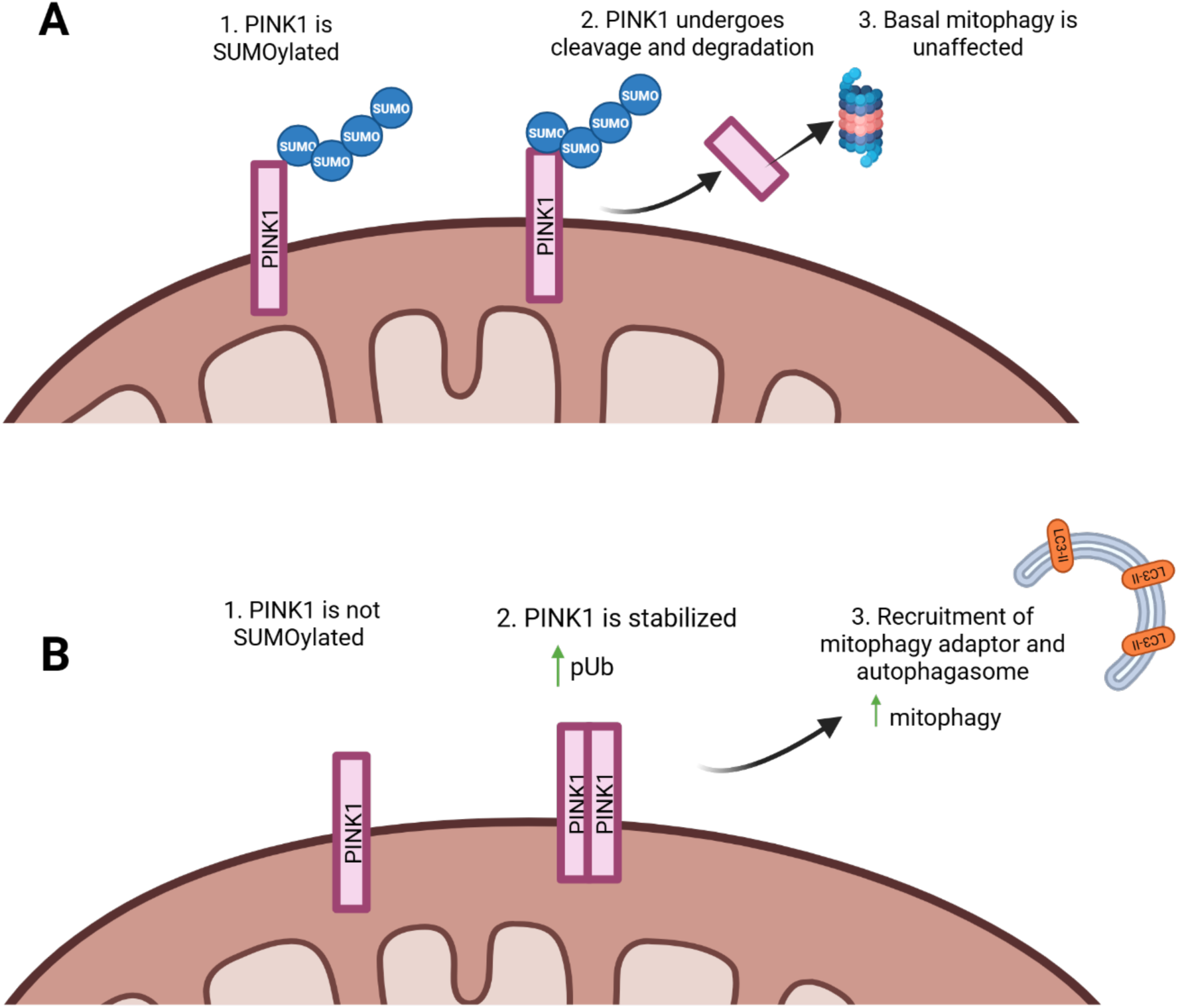
Proposed model of PINK1 SUMO-2/3-ylation and PINK1-mediated mitophagy. **(A)** When PINK1 is SUMO-2/3-ylated, it undergoes cleavage and is targeted for degradation, unaffecting basal mitophagy. **(B)** When PINK1 is not SUMO-2/3-ylated, PINK1 is stabilized, increasing levels of PINK1-mediated mitophagy. *Created in BioRender*.

DeSUMO-2/3-ylating PINK1 increased the ratio of LC3-II/LC3-I (Fig. 6E-6F), in agreement with our data from MAPL knockdown. Moreover, expression of PINK1-GFP in PINK1^-/-^ cells increased mitophagy (Fig. 6G-6H), and this effect was enhanced when PINK1-GFP was deSUMO-2/3-ylated (Fig. 6G-6H). Together, these results demonstrate that reducing PINK1 SUMO-2/3-ylation increases mitophagy.

## Discussion

Although most documented SUMO substrates are nuclear proteins, an extensive and increasing number of extranuclear SUMO substrates have been identified (Henley et al., 2018). In particular, multiple integral and associated mitochondrial proteins have emerged as SUMO2/3 substrates. These include dynamin-related protein 1 (Drp1), mitochondrial fission factor (MFF), mitochondrial fission 1 protein (Fis1), mitofusin-2 (Mfn2), and Fas-associated protein with Death Domain (FADD) (Braschi et al., 2009; Choi et al., 2017; Kim et al., 2021; Prudent et al., 2015; Seager et al., 2024; Waters et al., 2022). Here we validate PINK1 as a SUMO2/3 substrate, consistent with a previous proteomics screen to identify endogenous SUMO substrates, which identified PINK1 as a “hit”, but was not further investigated (Hendriks et al., 2018). After our data were posted on BioRxiv, another group reported that PINK1 can also be modified by SUMO-1 (J. Liu et al., 2025).

Several mitochondrial SUMO substrate proteins are involved in specific mitophagy pathways. For example, SUMO2/3-ylation of Fis1 at K149 is critical for its mitochondrial localisation, and SENP3-mediated deSUMO-2/3-ylation of Fis1 promotes mitophagy induced by the iron-chelator deferiprone (Waters et al., 2022). Mfn2 is SUMO2/3-ylated in response to CCCP or MG132 treatment, which promotes perinuclear aggregation of damaged mitochondria, and subsequent targeting for mitophagy (Kim et al., 2021).

### MAPL mediates PINK1 SUMO-2/3-ylation

We show MAPL-mediated SUMO2/3-ylation of PINK1 plays a key role in regulating mitophagy. MAPL is an OMM-localised SUMO and ubiquitin E3 ligase that can SUMOylate Drp1 and MFF (Prudent et al., 2015; Seager et al., 2024). MAPL knockdown significantly decreased SUMO2/3-ylated PINK1 and MAPL co-immunoprecipitated with PINK1, indicating that MAPL binds to and SUMOylates PINK1.

Three previous reports have suggested a link between MAPL and PINK1 in regulating mitophagy (Igarashi et al., 2020; Rojansky et al., 2016; Yun et al., 2014). Each study described a different effect, and none defined the role of SUMO-2/3-ylation. Briefly, using the anticancer drug gemcitabine, it was reported that MAPL stabilises PINK1 at the OMM to promote PINK1-dependent, Parkin-independent mitophagy (Igarashi et al., 2020). In contrast, using both *Drosophila* PINK1/Parkin and mouse cortical neuronal cultures Yun et al. reported that MAPL and PINK/Parkin act in parallel by regulating mitofusin ubiquitination and stability (Yun et al., 2014). The third study reported that MAPL and Parkin act downstream of PINK1 in parallel pathways to eliminate paternal mitochondria in mouse embryos (Rojansky et al., 2016).

Our results show that MAPL binds to, and promotes, PINK1 SUMO-2/3-ylation, providing new mechanistic understanding of the relationship between these key mitochondrial proteins.

### SUMO-2/3-ylation affects PINK1 stability

CCCP significantly decreased PINK1 SUMO2/3-ylation, and the cleavage-resistant PINK1 P95A mutant is less SUMO2/3-ylated than wild-type PINK1. These results suggest that SUMO-2/3-ylation prevents PINK1 stabilisation at the OMM and/or promotes PINK1 import and subsequent degradation. Furthermore, while reducing SUMO-2/3-ylation, both CCCP and the P95A PINK1 mutant increase PINK1 activity. MG132 significantly increases PINK1 SUMO-2/3-ylation, consistent with a proteomics study that identified PINK1 as a potential SUMO-2/3-ylation target in HEK293T cells stressed with MG132 (Hendriks et al., 2018).

In healthy mitochondria, PINK1 is constitutively turned over, and we show that, under basal conditions, this degradative process is regulated by PINK1 SUMO2/3-ylation.

### Site(s) of PINK1 SUMO-2/3-ylation

Despite exhaustive attempts using different approaches, we were unable to determine the sites of PINK1 SUMO2/3-ylation. Even a PINK1 construct in which all lysines were replaced with arginines was still SUMO-2/3-ylated. Recently, SUMO-2/3-ylation of two proteins, cofilin and p14ARF were shown to be lysine independent, with SUMO-2/3-ylation instead occurring at the N-terminal (El Motiam et al., 2025; Weng et al., 2023). We tested if this occurred for PINK1 by co-transfecting cells with PINK1-GFP and an irreversible N-terminal acetylase, Naa60, but this did not abolish PINK1 SUMO-2/3-ylation; thus, the exact locations of PINK1 SUMO2/3-ylation remain to be determined.

These results demonstrate that PINK1 undergoes non-canonical lysine-independent SUMO-2/3-ylation. Although unusual, there is established precedent for non-canonical ubiquitination of residues such as serines, cysteines, or threonines (Kelsall, 2022). Interestingly, the chemical reaction involving a nucleophilic acyl transfer during ubiquitination is similar to SUMO-2/3-ylation, providing a potential molecular basis for serine, threonine, or cysteine SUMO-2/3-ylation. Although outside the scope of the current study, these possibilities represent exciting future avenues of research.

### PINK1 SUMO-2/3-ylation and mitophagy

In agreement with previous reports (Cilenti et al., 2024; Li et al., 2015; Yuan et al., 2019), we show that MAPL knockdown increases the LC3-II/LC3-I ratio and p62 levels in WT cells. Crucially, these changes do not occur in PINK1^-/-^ HEK293T cells. Moreover, live imaging of cells transfected with mitoKeima showed increased mitophagy in response to MAPL knockdown in WT but not PINK1^-/-^ cells. These results indicate that MAPL plays a key role in the induction of PINK1-dependent mitophagy.

In our experiments we used global MAPL knockdown, so we cannot definitively exclude the possibility that the effects we observe on mitophagy could be accounted for by MAPL acting via proteins other than PINK1. Nonetheless, the lack of significant changes in mitophagy markers following MAPL knockdown in PINK1^-/-^ cells provides compelling evidence for a PINK1-dependent pathway. To more specifically interrogate PINK1 SUMO-2/3-ylation we targeted the SENP1 catalytic domain directly to PINK1-GFP. In agreement with the MAPL knockdown studies, these experiments showed that reducing PINK1 SUMO-2/3-ylation increases mitophagy.

### SUMO-1-ylation of PINK1?

A very recent study reported that mouse PINK1 is SUMO-1-ylated at K522 and K363 (corresponding to K364 and K523 in human PINK1) when overexpressed together with Ubc9 in SH-SY5Y or HEK-293 cells (J. Liu et al., 2025). Importantly, despite multiple attempts, and in the same experiments in which we reproducibly detected SUMO-2/3-ylation of human PINK1, we did not detect SUMO-1-ylation of PINK1. We also tested the K364R and K523R mutations individually (Fig. 2A), and a K364R/K523R double mutant (Fig. S3). As for our 21KR mutant, PINK1 SUMO2/3-ylation was still present (Fig 2B-2C, Fig. S2A-S2B).

In contrast to our findings for SUMO-2/3-ylation, Liu *et al*. suggest that SUMO-1-ylation stabilises PINK1 and increases stress-induced mitophagy (J. Liu et al., 2025). The reasons for this discrepancy are unclear, but there are substantive differences in the model systems and techniques used between the two studies. Thus, although we were unable to detect PINK1 SUMO-1-ylation in our experimental conditions, SUMO1 and SUMO2/3 may exert opposing actions. However, the discrepancy between the identification of the precise site(s) of PINK1 SUMO-2/3-ylation, irrespective of the SUMO paralogue, remains an outstanding question.

In conclusion, we show that SUMO2/3-ylation of PINK1 attenuates basal mitophagy. The removal of damaged mitochondria is critical for cellular health (Wang et al., 2023) so mitophagy must be stringently regulated to remove damaged but spare healthy mitochondria. We describe a novel pathway in which PINK1 SUMO2/3-ylation, mediated by MAPL, regulates basal mitophagy. These findings are important because understanding how PINK1 SUMO-2/3-ylation fits into the picture of basal mitophagy, and how it affects overall mitochondrial health and function will be essential for future therapeutic modulation of this pathway.

## Materials and Methods

### Reagents and antibodies

HEK293T cells were acquired from European Collection of Cell Cultures (ECACC). PINK1^-/-^ HEK293T cells (Abcam, ab266393) were a kind gift from Prof Ian Collinson (University of Bristol, UK). Dulbecco’s Modified Eagle’s Medium (D5796), fetal bovine serum (FBS), and poly-l-lysine (PLL), N-ethylmaleimide (E3876), Doxycycline (D3447), CCCP, and MG132 (47490) were purchased from Sigma. Sodium pyruvate (11360-070), Dulbecco’s Phosphate Buffered Saline (14200-059), L-glutamine (25030-024), 0.05% trypsin-EDTA and Dulbecco’s Modified Eagle Medium for live imaging experiments (A14430-01) was obtained from Gibco. Lipofectamine 2000 transfection reagent (11668-019) and sheep IgG Isotype Control (31243) were purchased from Invitrogen.

Complete protease inhibitor cocktail tablets and Bovine Serum Albumin Fraction V (10735094001) were obtained from Roche. Protein G Sepharose 4 Fast Flow beads (17-0618-01) were obtained from GE Healthcare Biosciences. ChromoTek GFP-trap and Spot-trap agarose beads were obtained from Proteintech. Enzymes and buffers used for cloning were obtained from New England Biolabs (NEB): Cut-smart buffer (B7204S), BamHI (R3136S), HindIII (R3104S), Dpn1 (R0176L), and Quick CIP (M0525S). Solution I ligase was obtained from Takara (6022-1).

For western blotting, a list of primary antibodies used in this study can be found in Table 1. HRP-conjugated secondary antibodies were used as follows: donkey anti-sheep IgG HRP (Abcam, Ab6900), goat anti-mouse IgG HRP (Sigma, A3682), rabbit anti-rat IgG HRP (Sigma, A5795), and goat anti-rabbit IgG HRP (Sigma, A6154).

**Table 1.**
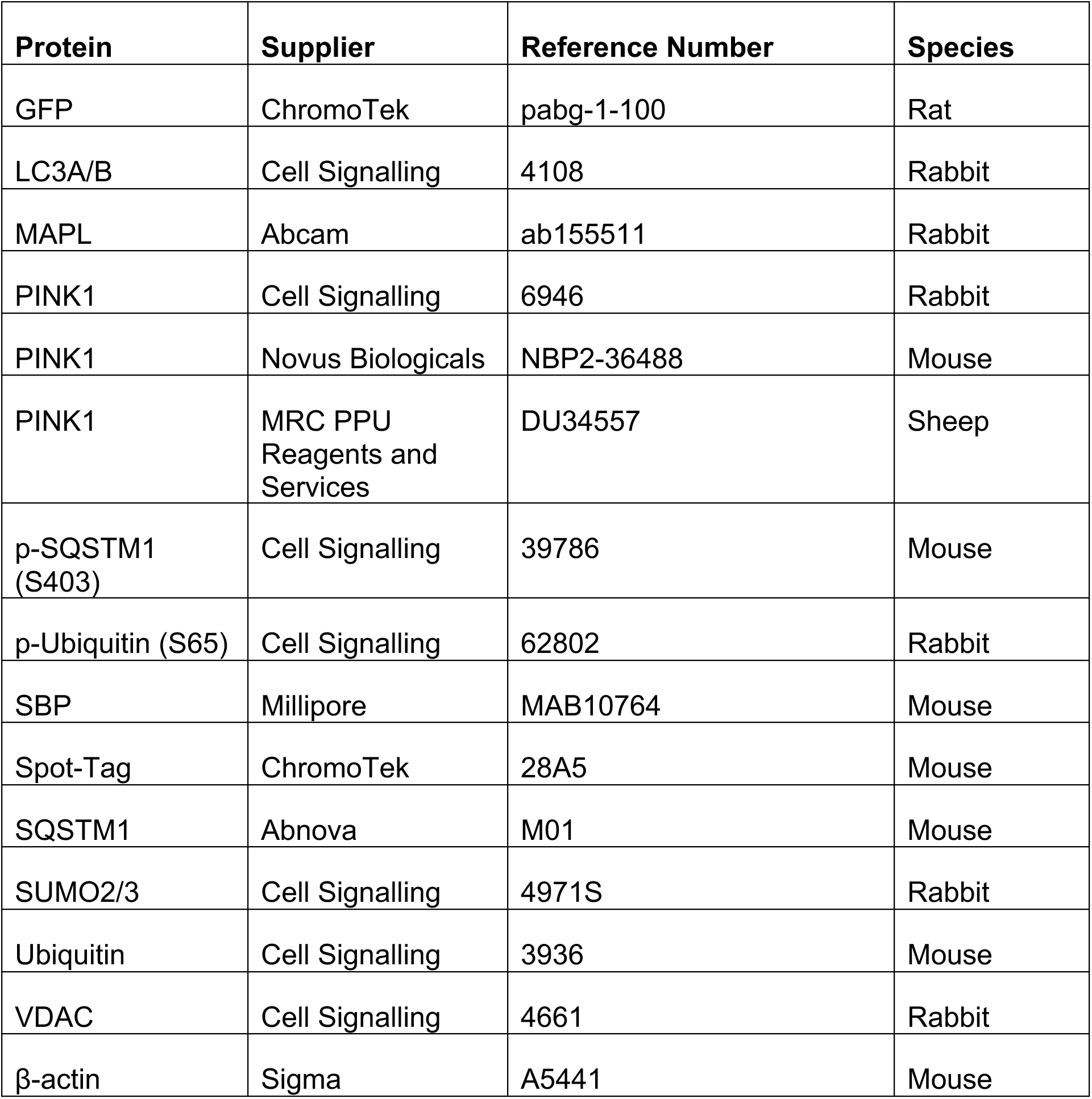
List of antibodies used for immunoblotting.

| **Protein** | **Supplier** | **Reference Number** | **Species** |
| --- | --- | --- | --- |
| GFP | ChromoTek | pabg-1-100 | Rat |
| LC3A/B | Cell Signalling | 4108 | Rabbit |
| MAPL | Abcam | ab155511 | Rabbit |
| PINK1 | Cell Signalling | 6946 | Rabbit |
| PINK1 | Novus Biologicals | NBP2-36488 | Mouse |
| PINK1 | MRC PPU<br>Reagents and<br>Services | DU34557 | Sheep |
| p-SQSTM1<br>(S403) | Cell Signalling | 39786 | Mouse |
| p-Ubiquitin (S65) | Cell Signalling | 62802 | Rabbit |
| SBP | Millipore | MAB10764 | Mouse |
| Spot-Tag | ChromoTek | 28A5 | Mouse |
| SQSTM1 | Abnova | M01 | Mouse |
| SUMO2/3 | Cell Signalling | 4971S | Rabbit |
| Ubiquitin | Cell Signalling | 3936 | Mouse |
| VDAC | Cell Signalling | 4661 | Rabbit |
| β-actin | Sigma | A5441 | Mouse |

### Plasmids and siRNA

For MAPL knockdown experiments, a SMART pool of siRNA targeting MAPL (Dharmacon, L-007062-00-0005) or a non-targeting pool (Dharmacon (D-001810-10-05) was used. To generate PINK1-GFP and mutant plasmids, RNA was extracted from HEK293T cells using the Qiagen RNAeasy mini kit, cDNA was made from this RNA using Thermo RevertAid First Strand Kit, and human PINK1 sequence was amplified from this cDNA by PCR using the following primers: Fwd: CACAAGCTTGCCACCATGGCGGTGCGACAGGCGCTGGGC, Rev: GTGGGATCCTCCAGGGCTGCCCTCCATGAGCAGAG. The PCR product (insert) and vector (peGFP-N1) was then digested using BamH1 and HindIII, ligated at a 3:1 ratio of plasmid:insert, and transformed into DH5α bacteria. The GFP tag was added to the C-terminus of PINK1 since it has previously been shown that N-terminally tagged PINK1 is trapped in the mitochondria, possibly due to N-terminal cleavage of PINK1 (Beilina et al., 2005). For site-directed mutagenesis and PINK1-SPOT, mutants were made using WT PINK1-GFP as the template and mutagenesis primers were comprised of 21 base pairs before and after the desired mutation. 1 µL of Dpn1 was added to this PCR product for 1 hour at 37°C before being purified using GeneJet Gel Extraction kit (#K0691) and transformed into DH5α bacteria.

The mitochondria-targeted Keima probe (pMitophagy Keima Red mPark2, from MBL (AM-V0259M)) was a kind gift from Prof Ian Collinson (University of Bristol, UK). GNb and GNb-SENP1 were modified from a GFP nanobody-NLS-catalytic SENP1 plasmid from Prof Ron Hay (University of Dundee, UK) as described in (Mojsa et al., 2020). Briefly, GNb was produced by introducing a stop codon after the GNb coding sequence, and the GNb-SENP1 construct was modified to remove the nuclear localisation sequence and favor cytosolic expression of the SENP1-nanobody fusion protein.

SBP-tagged SENP1, 3, 5, 6 and 7 constructs were produced by amplifying the corresponding coding regions from rat neuronal cDNA and cloning into a variant of the plasmid pcDNA3.1 containing an N-terminal SBP tag.

### Cell culture and transfection

HEK293T cells were grown in complete media (Dulbecco’s Modified Eagle Medium (DMEM) with 10% fetal bovine serum (FBS), penicillin (1000U)), and streptomycin (0.1mg). Cells were passaged regularly in T75 flasks and used for experiments between passages 2 and 20. For immunoprecipitation experiments, approximately 1.75 million HEK293T cells were plated into 60 mm dishes and transfected when ∼60-70% confluent. In plain DMEM (without FBS, penicillin, or streptomycin), 2.5 µg of DNA (for a double transfection) or 5 µg of DNA (for a single transfection of DNA) was added along with lipofectamine 2000 (Thermo Fisher, 1.5 µL/µg of DNA). For siRNA-mediated knockdown, JetPrime transfection reagent (Polyplus) was used. Similarly, siRNA was added to JetPrime buffer, along with JetPrime transfection reagent (2 µL/µg). This transfection mix was kept at room temperature for 30 minutes and then added dropwise onto cells.

### Immunoblotting

Total cell lysates or immunoprecipitation samples were run on 10% or 15% SDS-PAGE gels and for 90 minutes at 120V. Proteins were transferred onto PVDF membranes at 400mA for 90-120 minutes. PVDF membranes were blocked with 5% milk (low-fat milk powder in PBST) or 5% BSA (bovine serum albumin in PBST) for one hour. Membranes were incubated with primary antibodies (Table 1) for one hour at room temperature or overnight at 4°C, and HRP-conjugated secondary antibody for one hour at room temperature. Membranes were placed directly in 1 mL of HRP substrates and developed on the Odyssey Fc LiCor system. Bands were quantified using Image Studio.

### Immunoprecipitation and Co-immunoprecipitation

48-72 hours post transfection, cells were lysed in lysis buffer (150 mM NaCl, 50 mM Tris, 1% Triton, protease inhibitors, 20 mM NEM, and 0.1%-2% SDS), sonicated, and spun down for 20 minutes at 16,100g. The soluble fraction was then added to GFP-Trap or Spot-Trap beads (ChromoTek) and gently rotated for 1 hour at 4°C to immunoprecipitate GFP-tagged proteins. Beads were washed 3X with wash buffer (lysis buffer without NEM, protease inhibitors, or SDS). Sample buffer (125 mM Tris pH 6.4, 4% SDS, 10% glycerol, 0.004% bromophenol blue, 10% β-mercaptoethanol) was added to the beads and boiled for 10 minutes at 95°C. Before immunoprecipitation, 6% of total cell lysate was taken for input, added to sample buffer, and boiled for 10 minutes at 95°C. For co-immunoprecipitation experiments, cells were lysed in (20 mM Tris, 137 mM NaCl, 2 mM Na_4_P_2_O_7_, 2 mM EDTA, 1% Triton, 25 mM β-glycerophosphate, 10% glycerol, protease inhibitors, and 20 mM NEM) and samples were not sonicated.

### Mitochondrial Isolation and Endogenous Immunoprecipitation of PINK1

Confluent 150 mm dishes of HEK293T cells were detached using trypsin, centrifuged 1500 rpm 3 mins, washed once with 1XPBS, pelleted, and frozen at -80° overnight. Cells were then thawed on ice, and mitochondria were isolated as per the manufacturer’s protocol (Mitochondrial Isolation Kit for Cultured Cells, Abcam, ab110170). (Note: 20 mM NEM was added to each of the reagents (A, B, and C) provided in the kit to inhibit sentrin-specific protease activity). Mitochondrial pellet retrieved from isolation protocol was then resuspended in 1x TGS buffer (0.1M Tris-HCl, 0.15M NaCl, 10% glycerol, 0.5% glyco-diosgenin, 20 mM NEM, and protease inhibitors), incubated on ice for 15 minutes, and centrifuged at 13,200 rpm for 10 minutes at 4°C. Lysates were pre-cleared with 30 μL of protein G-sepharose beads for 2 hours. 10% of the pre-cleared samples were taken for total lysate, added to sample buffer, and boiled 95°C for 10 minutes. Sheep anti-PINK1 (S085D) or control sheep-IgG (Thermo, #31243) was incubated with pre-cleared lysate overnight. Samples were then incubated with 20 μL of protein G beads for 1 hour. Beads were washed 3X, added to sample buffer, and boiled 95°C for 10 minutes.

### In-vitro deSUMO-2/3-ylation and deubiquitination assay

PINK1-GFP was transfected in HEK293T cells and immunoprecipitated using GFP-trap beads as described above. Beads were washed three times, and then subsequently treated with constitutively active SENP1 (100 nM, purified in-house as described in (Rocca et al., 2017)) or active USP2 (500 nM, Biotechne, E-504) for 2 hours at 37°C. 2X laemmli buffer was added to the samples, boiled 95°C for 10 minutes, run on SDS-PAGE, and immunoblotted for SUMO2/3 and ubiquitin.

### mKeima-Red mitophagy assay

The mitochondrial-targeted mKeima probe (pMitophagy Keima-Red mPark2) has a bimodal excitation spectrum that is sensitive to pH, with an excitation wavelength of 405 nm around a pH of 8, and 561 nm around a pH of 4. The ratio of these two fluorescence values can be used as measure of mitophagy, with greater fluorescence in 561 nm excitation channel denoting more mitophagy. 35 mm glass bottom dishes (Greiner) were coated with poly-L-lysine for at least one hour in 37°C to increase cell adhesion. 250,000 cells were seeded in each dish and transfected the following day. 36-48 hours after transfection, cells were imaged live using the SpinSR microscope. At least one hour before imaging, cell media was changed to “extra-buffering” media without phenol red (DMEM (A14430-01), 10% FBS, 2mM L-glutamine, 4.5 g/L glucose, 1mM sodium pyruvate, 40 mM HEPES pH 7.4). Cells were imaged live on the Olympus IXplore SpinSR system (Yokogawa CSU-W1 SoRa spinning disk). Images were acquired using a 60X oil immersion lens (1.5 numerical aperture, 0.11 mm working distance). During image acquisition, glass dishes were kept in a 37°C chamber with constant CO2 supply. All acquisition settings (laser power, gain, exposure time, etc.) were kept constant across all repeats and conditions within each experiment. Cells were excited using 405 nm, 488nm, and 561 nm lasers and z-stacks of 10 µm range with 0.2 µm step size was obtained. To quantify percent of mitophagy (“mitophagy index”), the fluorescence in the 561 nm channel was divided by the total fluorescence (a sum of both 405 and 561 nm).

### Statistical analysis

A one-sample t-test (where the control was set to 100%) or as appropriate, a one-way or two-way ANOVA followed by Tukey’s post-hoc test was conducted to determine significance. For comparisons with more than two groups, all data was normalized to the average of control condition values across all repeats. For imaging experiments, within one biological repeat, all values were normalized to the average of the control condition. Outliers were removed using the ROUT method (Q= 1%). D’Agostino & Pearson test was used to test normality of data, and Kruskal-Wallis test (followed by Dunn’s post-hoc test) was conducted to test significance. All data are shown as mean ± SEM. Graphs and statistical tests were done on GraphPad Prism.

## Acknowledgements

The authors thank Prof Ian Collinson (University of Bristol, UK) for providing the mitochondria-targeted Keima probe and PINK1-/- HEK293T cells. We are grateful to Prof Ron Hay (University of Dundee, UK) for the GFP nanobody-NLS-catalytic SENP1 plasmid. The authors also gratefully acknowledge the Wolfson Bioimaging Facility (University of Bristol, UK) for their support and assistance in this work.

---

**Figure S1.**
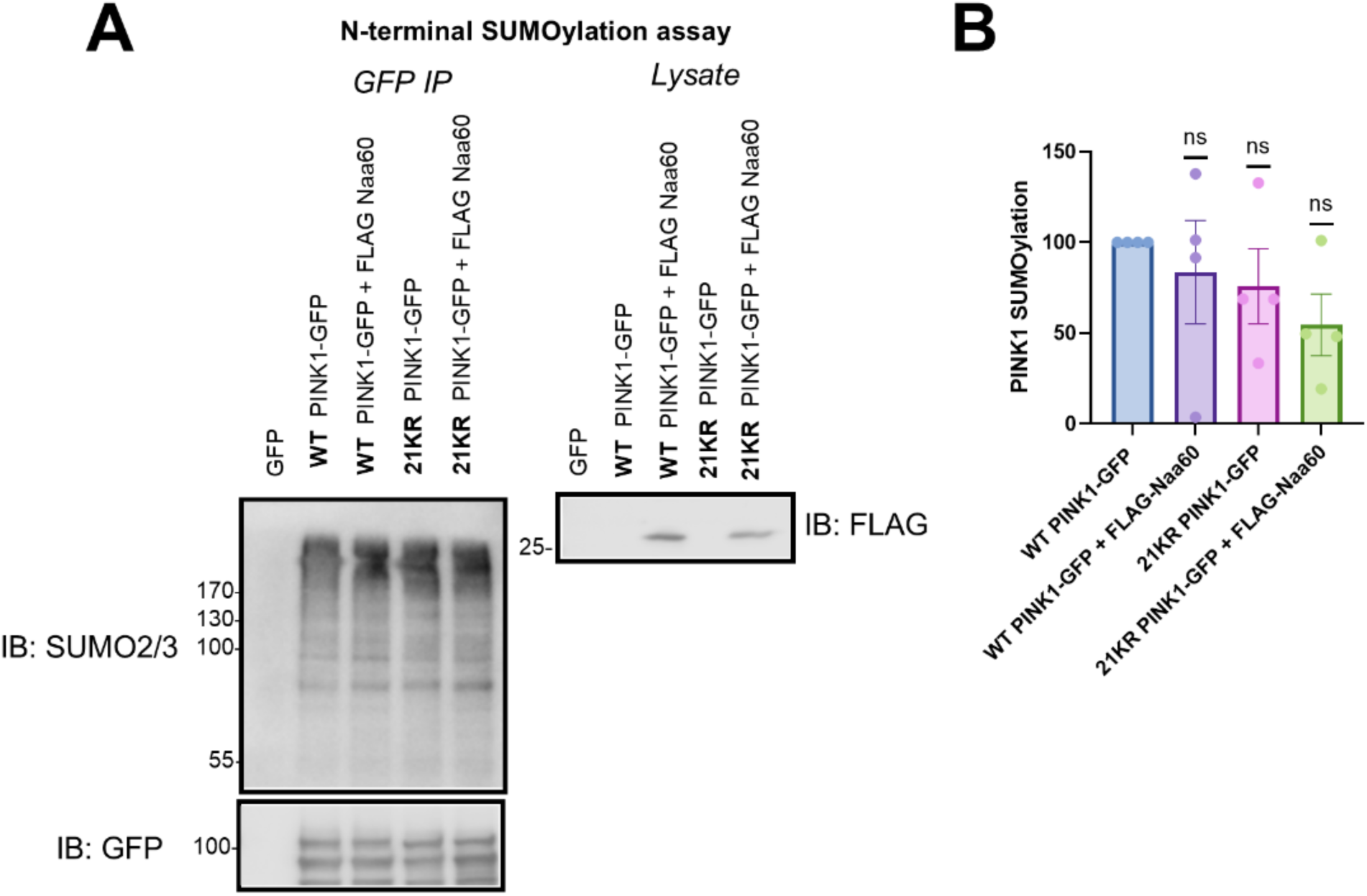
PINK1 is likely not SUMO-2/3-ylated at the N-terminal end. HEK cells were transfected with WT/21KR PINK1-GFP along with FLAG-Naa60 for 48 hours. Cells were lysed in buffer containing 2% SDS, and PINK1 was immunoprecipitated using GFP trap beads. Samples were immunoblotted for SUMO2/3, PINK1, and FLAG. PINK1 SUMO-2/3-ylation levels were quantified and compared to WT PINK1-GFP (SUMO2/3/GFP) (n=4, one-sample t-test)

**Figure S2.**
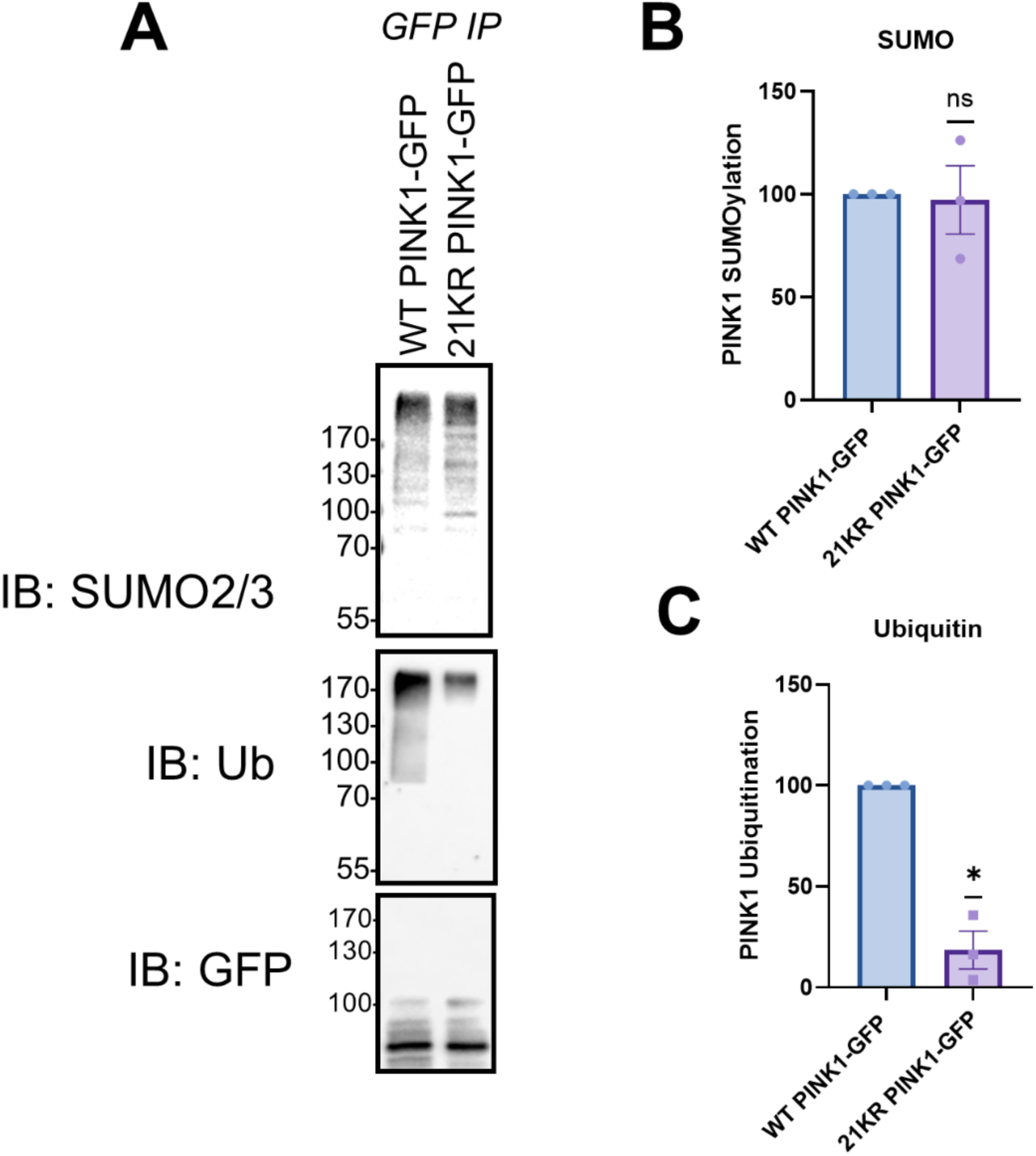
SUMO-2/3-ylation and ubiquitination levels of 21KR PINK1-GFP. HEK293T cells were transfected with PINK1-GFP or 21KR PINK1-GFP and PINK1-GFP was immunoprecipitated. **(A)** Representative immunoblot of SUMO2/3 and ubiquitin for immunoprecipitated samples **(B)** Quantification of SUMO2/3 smear on PINK1-GFP immunoprecipitation (SUMO2/3/GFP normalized to control (WT PINK1-GFP) **(C)** Quantification of PINK1-GFP ubiquitination (SUMO2/3/GFP normalized to control (WT PINK1-GFP) (n=3, one-sample t-test, *p<0.05)

**Figure S3.**
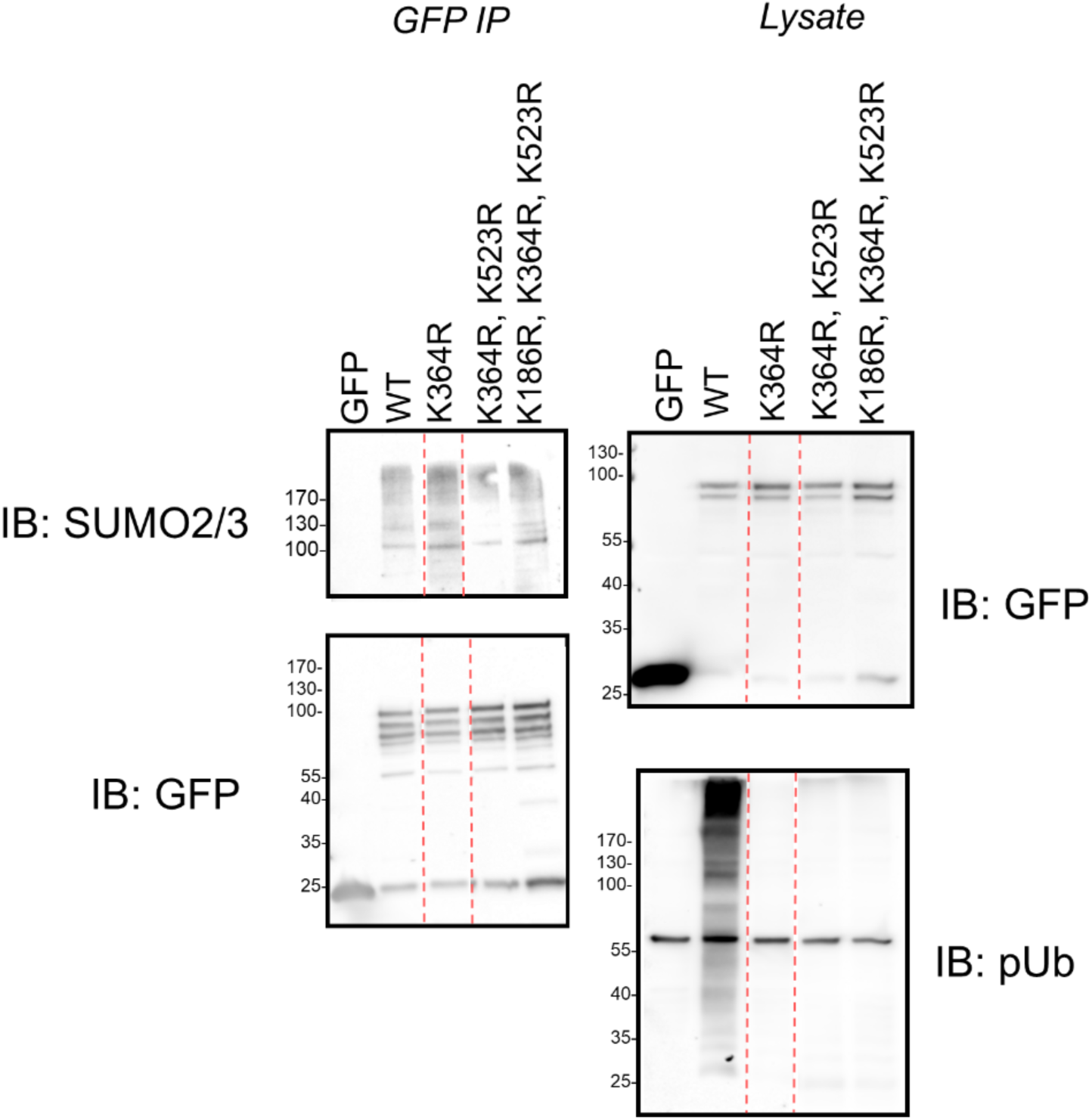
PINK1 is not SUMO-2/3-ylated at proposed SUMO1 sites, K364/K523. HEK cells were transfected with WT, K364, K364/K523R, or K186R/K364R/K523R PINK1-GFP. Cells were lysed in buffer containing 2% SDS, and PINK1 was immunoprecipitated using GFP trap beads. Samples were immunoblotted for SUMO2/3, GFP, or phospho-ubiquitin.

